# Volumetric Segmentation of Cell Cycle Markers in Confocal Images

**DOI:** 10.1101/707257

**Authors:** Faraz Ahmad Khan, Ute Voß, Michael P Pound, Andrew P French

## Abstract

Understanding plant growth processes is important for many aspects of biology and food security. Automating the observations of plant development – a process referred to as plant phenotyping – is increasingly important in the plant sciences, and is often a bottleneck. Automated tools are required to analyse the data in images depicting plant growth. In this paper, a deep learning approach is developed to locate fluorescent markers in 3D timeseries microscopy images. The approach is not dependant on marker morphology; only simple 3D point location annotations are required for training. The approach is evaluated on an unseen timeseries comprising several volumes, capturing growth of plants. Results are encouraging, with an average recall of 0.97 and average F-score of 0.78, despite only a very limited number of simple training annotations. In addition, an in-depth analysis of appropriate loss functions is conducted. To accompany [the finally-published] paper we are releasing the 4D point annotation tool used to generate the annotations, in the form of a plugin for the popular ImageJ (Fiji) software. Network models will be released online.

## II. INTRODUCTION

Understanding plant growth is becoming increasingly important, as the ability to feed an increasing population is heavily dependent on successful and efficient crop production. Growth of plants occurs as an interaction between two key activities: cell division and cell expansion. In developmental biology, quantifying cell divisions is an important measure to estimate and compare organ or tissue growth between different genotypes, or to compare different growth conditions. In plants, the root meristem is located at the root tip and overall root growth is achieved by the sum of generating new cells by cell division and their subsequent elongation. Therefore quantifying cell division is important when determining root growth dynamics. Being able to analyse these events is critical in many plant science experiments. Modern microscopy methods such as confocal or light sheet microscopy allow a biologist to see inside a plant root, at a cellular scale, and even over time. Analysis of these datasets for morphological changes remains challenging, as they often comprise large, possibly 4D image data. Certain fluorescent markers can be used to help visualise events such as cell division, but nevertheless, finding and counting the markers in these datasets remains a labour intensive task.

3D confocal microscopy is an imaging technology that allows us to see inside biological samples. Fluorescent markers or dyes are excited by laser light, allowing for clear labelling of particular structures within an organism. Volumetric image datasets can therefore be produced allowing inspection at the cellular scale. If timelapse imaging is used, such volumes can be collected at regular timepoints. The resulting “4D” datasets are large, and due to their dimensionality are hard to inspect by hand. Labelling regions in 3D data is often reduced to labelling 2D slices, or drawing complex surfaces. Both methods are time consuming and often frustrating to perform in practice. This challenge is the motivation behind the work presented here: developing a deep machine learning approach to segment and locate fluorescing nuclei markers inside an Arabidopsis root. Furthermore, the markers for the plant cells used in this paper provide a unique challenge, as the structures are small and sparse, and ill-defined, and are marked by the same colour as other distracting features in the image such as the cell walls.

## III. RELATED WORK

Segmentation of nuclei in 3D has presented image analysis with a challenge for many years. Indeed, labelling and segmenting cells and sub-cellular features such as nuclei has been an active research area for decades [1]. There is a need across many biological disciplines to locate, count and segment nuclei and nuclear markers for quantification in many types of experiments [2]. Until recently, handling the challenges of nuclei segmentation, especially in close proximity, has traditionally been accomplished by concatenating a pipeline of various analysis techniques together to separate the features in the image (e.g. [3], [2]). Now, machine learning-based methods are providing much more reliable results across a wider range of datasets without the need to use many individual processing steps. Deep machine learning in particular has begun to challenge preconceptions about how to accomplish image analysis tasks. Here, we will describe a deep machine learning approach to segment cell-cycle nuclear markers in 4D datasets, thereby developing an AI-based solution to a problem which has remained a challenge for many years, and will only become more important to solve as we are able to collect more, and larger, datasets.

It is certain that over coming years we will see a trend for traditional image analysis and processing pipelines to be replaced or at least supported by dedicated deep learning models. Some existing popular tools for cell image analysis, such as CellProfiler [4] have recently begun to support the ability to load in developed deep learning models to form part of their analysis suite. Beyond segmentation, other applications of deep learning with reference to microscopy include approaches to generate super-resolution images – software-enhanced images creating clarity beyond the physical limits of the microscope system in use [5]. 3D segmentation using deep learning presents particular challenges. Most deep learning to date has focused on 2D data, requiring significantly fewer computational resources due to the size of the datasets in use. Care must be taken when developing fully volumetric approaches, as the limits of even modern computer systems are easily reached.

The work here focuses on plant cells, but 3D microscopy segmentation using deep learning is an active area of research in many domains. Recent work has developed a deep network approach to analysing neurites in brain EM images [6]. Convolutional neural networks have also been used to segment larval zebrafish intestines in light sheet images [7]. This was found to be as accurate as human experts, and outperformed other non-deep machine learning approaches in common use, such as random forest and support vector machines. MRI (Magnetic Resonance Imaging), another 3D imaging technique, has also received attention from the deep learning community, applications of which include brain scan analysis [8], and diagnostics for knee imaging [9].

When segmenting regions of interest in 3D several deep learning architectures may be used, but all have a common structure. Key to any form of semantic segmentation is to enable an efficient upscaling in the network so that a high-quality, pixel-wise map can be produced for the segmentation labels. Convolutions are used to reduce the initial image dimension down to a spatially-reduced set of dense, high-level features. These are then upsampled back into pixel values at a scale normally equivalent to the input image. Of course, as spatial information is lost in the central part of the architecture, techniques have been developed to maintain spatial information throughout this process. These include skip layers in the Fully Convolutional Network (e.g. [10]) or in the case of U-Net, many feature channels are additionally used in the upscaling path to preserve features [11]. This latter 2D approach is expanded to handle volumetric data with 3D U-Net [12]. In particular, the authors recognise the challenge of annotating 3D data to provide as training instances, so propose a 2D annotation approach. The network is envisioned to both fill in sparse 2D annotations to produce a 3D dataset segmentation, and to generalise to new unannotated datasets. Both applications are relevant to this work, so a 3D encoder-decoder architecture based on a modified U-Net will be the base architecture used here.

The rest of this paper is organised as follows. In the following section we describe the nature of the microscopy involved and the dataset details. This is followed by an explanation of the deep learning architecture developed, and the approach taken for data augmentation. Choice of loss function is a critical aspect of the design, and different loss functions are explored at the end of the Methodology section. We then present quantitative and image-based results of the developed deep learning approach on unseen data, and the results are then explored in the discussion. This includes an examination of the failure cases of the network, and proposed methods of handling such errors in the future.

## IV. METHODOLOGY

### A. The Confocal Dataset

The confocal dataset used here is composed of five 3D time-lapse sequences depicting five g rowing A rabidopsis thaliana roots. To capture images of cell divisions, confocal microscopy time course datasets have been generated. Capturing cell division events is challenging, as the divisions (representing the emergence of new cell walls) happens relatively quickly, so this process is likely to occur between image capture points. Therefore, identification of dividing cells is facilitated here by the use of specialist marker lines, in which fluorescent cell division-specific p roteins a llow t he v isualisation o f dividing cells. In particular we made use of the Arabidopsis thaliana line CYCB1;1:: CYCB1;1-GFP [14] which marks nuclei of dividing cells. Imaging was carried out on a Leica SP8 Confocal microscope. The fluorescing markers can be seen in the captured image as bright spots in Figure 2(a, c), for example.

To visualise and quantify this cell division in Arabidopsis, we imaged growing roots of 5 day old seedlings (grown at 22 degrees Celsius in 12h light 12h dark cycles on 0.5 MS, pH 5.8) for up to 3 hours, every 10 – 30 min. To train the software, every cell division event was manually annotated using “Orthogonal Pixels” in Fiji [13]. This is a custom written plugin which alleviates the challenge of labelling broadly-spherical 3D structures as points in volumetric data. The problem with capturing such data is that it can be a challenge to identify where the centre of such a structure lies, and additionally in identifying if a structure has already been labelled on a neighbouring slice. This tools allows efficient capture of point annotations in 3D or 4D by representing 3D clicks visually as spheres. At each stack spheres representing clicks in nearby slices are also visualised, reducing the chance of annotating the same object twice (see Figure 1). This plugin is available as a supplemental file to [the final] paper.

**Figure 1:**
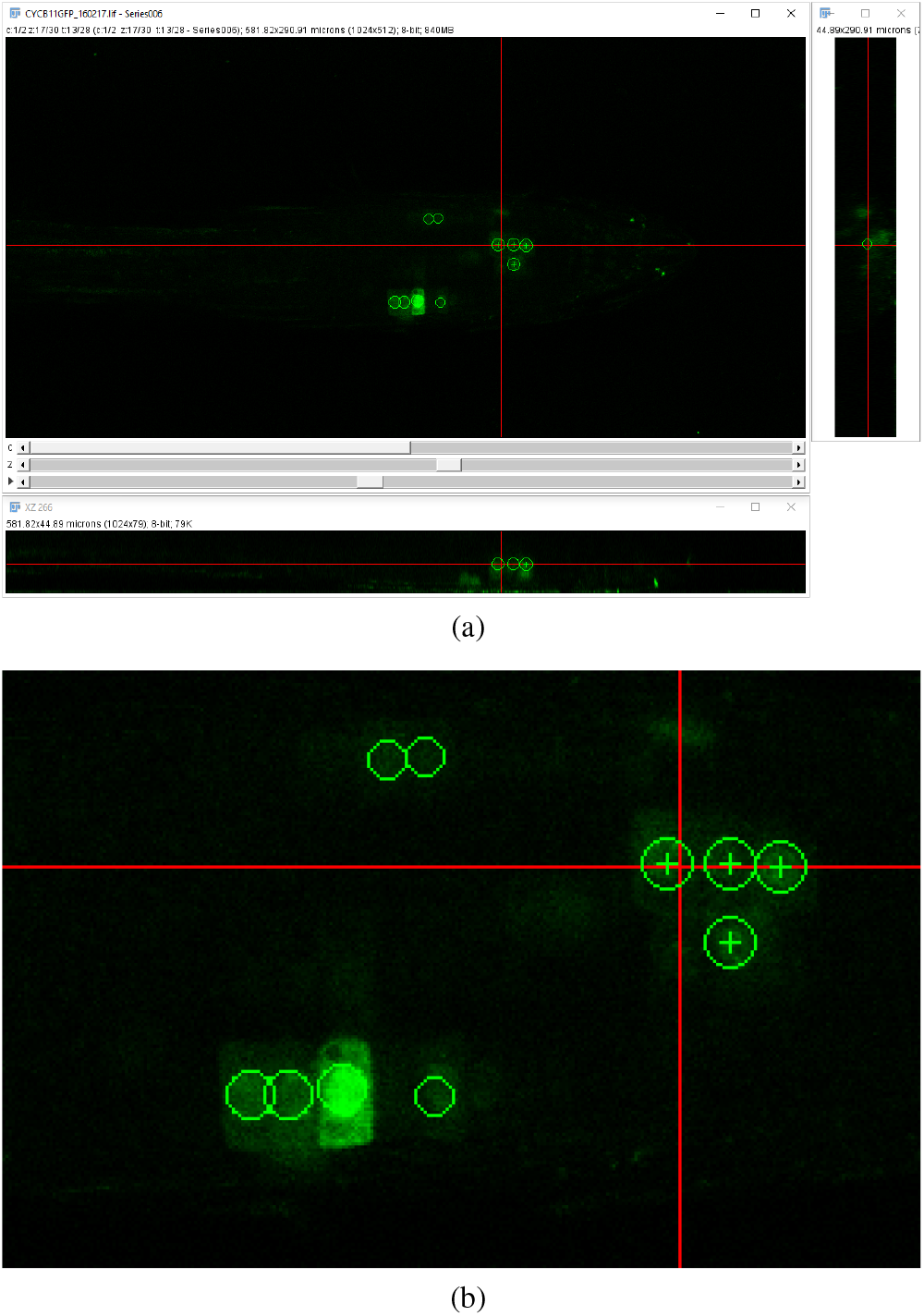
(a) Annotation of cell division events using the orthogonal pixels plugin in Fiji [13]. (b) Zoom of (a). Note that point annotations are visualised as spheres – circles are seen in each of the orthogonal views. This makes labelling of 3D data in individual views easier for the annotator, and prevents common errors, such as double labelling of single structures. Crosses indicate the centre of the sphere, indicating the plane on which they were placed

**Figure 2:**
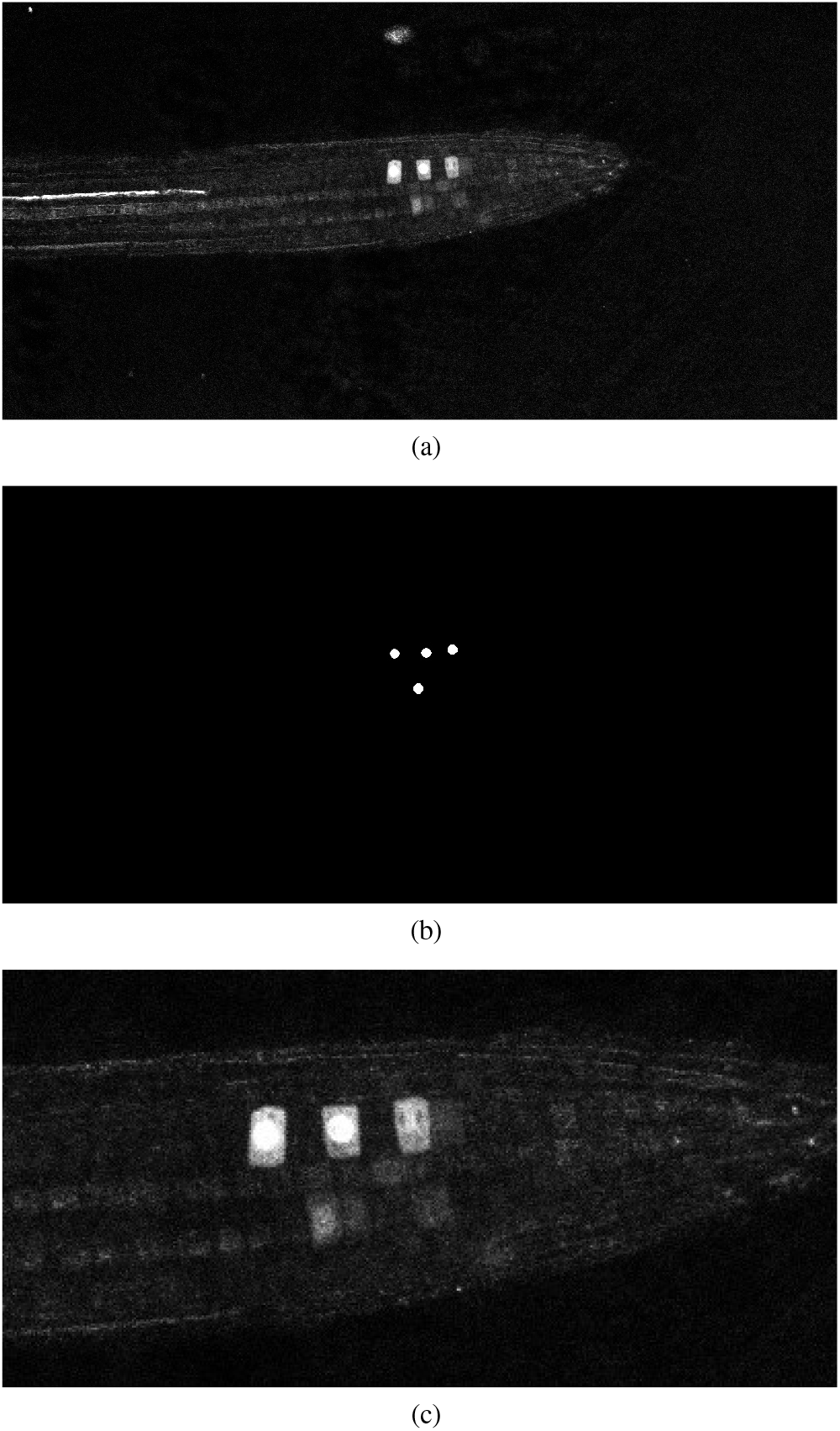
(a) Shows a 2D slice from a 3D confocal volume, and its corresponding ground truth can be seen in (b). (c) shows zoomed details of panel (a), highlighting the fluorescing cell-cycle markers, as well as bleed-through marker fluorescence and noise.

In total, a little over 1000 cell division events were annotated. To enable efficient labelling, all such events in the dataset have been annotated by a single expert, stored as *x, y, z* and *t* coordinates representing the centre of each nucleus in 3D space and at a point in time. These have been manually localised in the datasets by navigating to the *x, y, z, t* position within the data using the plugin, and adding an annotation with the space bar. Annotations are then stored as a CSV text file alongside the volumetric image data. Note that although individual time points are used here, we anticipate the approach being deployed on data captured over longer periods and more frequent time points using technology such as light sheet microscopy. A comparison of confocal microscopy and lightsheet microscopy and the challenges of imaging with plants can be found in [15]. Importantly, the data resolves to ultimately the same format as used here, both being volumetric RGB image stacks, exported from the manufacturer’s microscopy software as uncompressed TIFF files.

The point *x, y, z* coordinates at a specific time need further processing in order to effectively train a semantic segmentation network. Our point labels must be expanded to encompass a volume of space, which is centred on the feature we are interested in. To this end, in order to generate a ground truth volume for training purposes, a 3D Gaussian function is used to generate a region of interest at every *x, y, z* coordinate. This provides a contextual volume of space to the network centered on the area of interest. As the marker is concentrated in the nucleus with some bleed into the surround cell space and surrounding space, this is an appropriate representation.

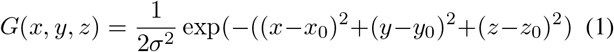

Here, *x*_0_, *y*_0_, *z*_0_ represent the centre of the are of interest, and *σ* represents the standard deviation of the Gaussian curve.

The standard deviation effectively defines an area of interest centred on the click locations. Depending on the loss function in use, this Gaussian can be used to define contextual interest which drops off away from the centre, or a threshold can be applied to effectively produce fixed radius spheres around the annotation points. This latter approach produces a binary 3D volume where the areas of interest (nuclei in this case) are represented in a sphere defined by logical 1’s and anything other than the nuclei are represented by logical 0’s. A 2D slice from a 3D volume can be seen in Figure 2 (a), while its generated ground truth can be seen in Figure 2 (b) where a 3D Gaussian is generated at every point of interest from a ground truth coordinate file accompanying the image data. In the case of Figure 2 (b), this is then thresholded at a fixed distance to produce a spherical region.

### B. Network Architecture

As mentioned above, our network utilizes the encoder-decoder approach based on a modified version of the 3D U-Net presented in[12]. Figure 3 shows an outline of the network architecture used. As can be seen, the network path is composed of 4 encoder blocks, 3 decoder blocks and a final convolution block. Each encoder block consists of two consecutive convolution blocks followed by a MaxPool layer. Each convolution block is composed of a convolution layer, followed by a Rectified Linear Unit (ReLU) [16] layer and a batch normalization layer. Each decoder block is composed of an upsampling layer that reverses the process of MaxPool performed at the encoder stage. This is followed by two consecutive convolution blocks, similar in structure to those used at the encoder stage. The last convolution block performs a single [1 × 1 × 1] convolution that reduces the number of output channels to the number of classes, so in our case, for a single marker, the final layer will reduce the output to a single channel. Finally, an additional sigmoid layer is used to normalize the network output so that the unnormalized logits can be fit to the target. Cropping is avoided in the decoder path by using padded convolutions, this ensures that the output resolution of the convolution layer is the same as that of the input. The resulting network architecture has a total of 59,329,153 learn-able parameters.

**Figure 3:**
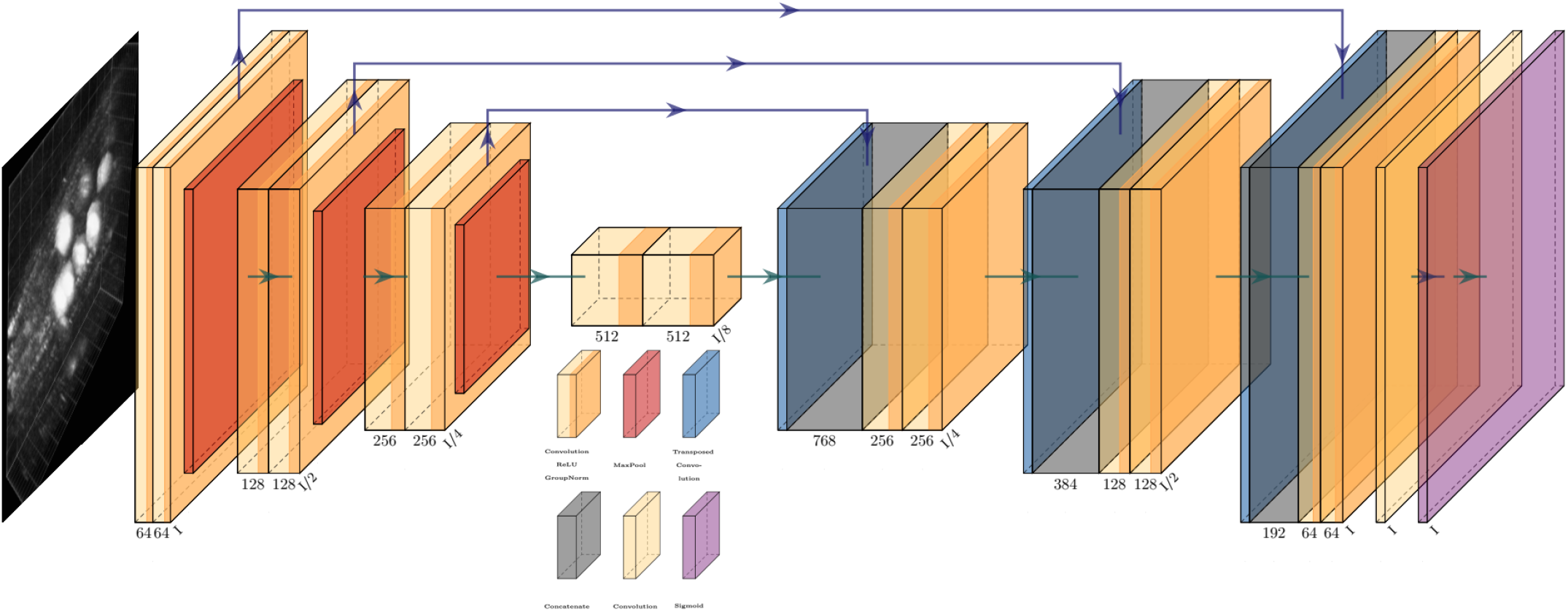
An overview of the network architecture used. Key to blocks – yellow : convolution; red : max pooling; blue: transposed convolution; grey: concatenation; purple: sigmoid.

In our case, upsampling via transposed convolution was favored over a pre-defined interpolation approach because it was observed that in cases of severe class imbalance, a learned upsampling approach performed better than simple interpolation. One possible reason for this being that the input to the upsampling layer is rather coarse, and straightforward interpolation is too simple a process to capture subtle features. Due to this, small objects or regions (such as the marker being segmented here) are often overlooked or misclassified [17]. This transposed convolution process reconstructs the previous spatial resolution and follows it with a convolution operation [18].

The confocal volumes used here are of the dimensions 1024 × 512 × 32 in the *x*, *y* and *z* directions, respectively. This is too large a volume of data to be input directly into the network as is, since at such sizes GPU memory becomes a limiting factor. This is a common problem when working with 3D data in deep learning methodologies. Instead of feeding in all data at once, we used an overlapping patch-based approach to train the network, with the full volume divided into patches of size 128 × 128 × 20. Patches are overlapped rather than tiled to ensure that the chance of a feature of interest (a nucleus in this case) occurring at the edge of a patch and being cropped is minimised. A 25% overlap between patches in all directions is used.

### C. Data Augmentation

We observed fairly consistent appearance between samples across time series and images, and so felt that extensive and complex augmentations such as elastic deformations, skewing and random angle rotations were not necessary during training. We first normalised the dataset samples to a zero mean and unit variance, followed by standard augmentation such as random flipping of the volume over the x-axis and random 90 degree rotations, representing different presentations of the root structure to the microscope. Given more complex datasets or perhaps different presentation to the microscope, additional augmentation of the data could be incorporated.

### D. Loss Function

Successful segmentation is made possible not only by the architecture of the network used, but also by the use of an effective loss function for the task at hand. The choice of loss function is even more important where there exists a severe class imbalance between the background and foreground objects. This is the case here, where the markers used represent a small proportion of the root, which itself occupies only a fraction of the complete image volume.

Recently in literature, Weighted Cross Entropy (WCE) [11], Binary Cross Entropy (BCE), Dice [19] and Generalized Dice Loss (GDL) [20] have been used as loss functions to segment data when there is a high imbalance between the foreground and background pixels. To find the best performing loss function for our dataset, separate instances of our network were trained using each of these loss functions, and the validation results produced by early stages of training were compared. Figure 4 shows a comparison of the network output using different loss functions. As can be observed, for the binary segmentation of such an imbalanced dataset Dice and GDL appear to perform better than WCE and BCE. In the end a decision was made in favour of GDL as it takes into consideration and utilizes the class-rebalancing property of the Dice overlap [20].

**Figure 4:**
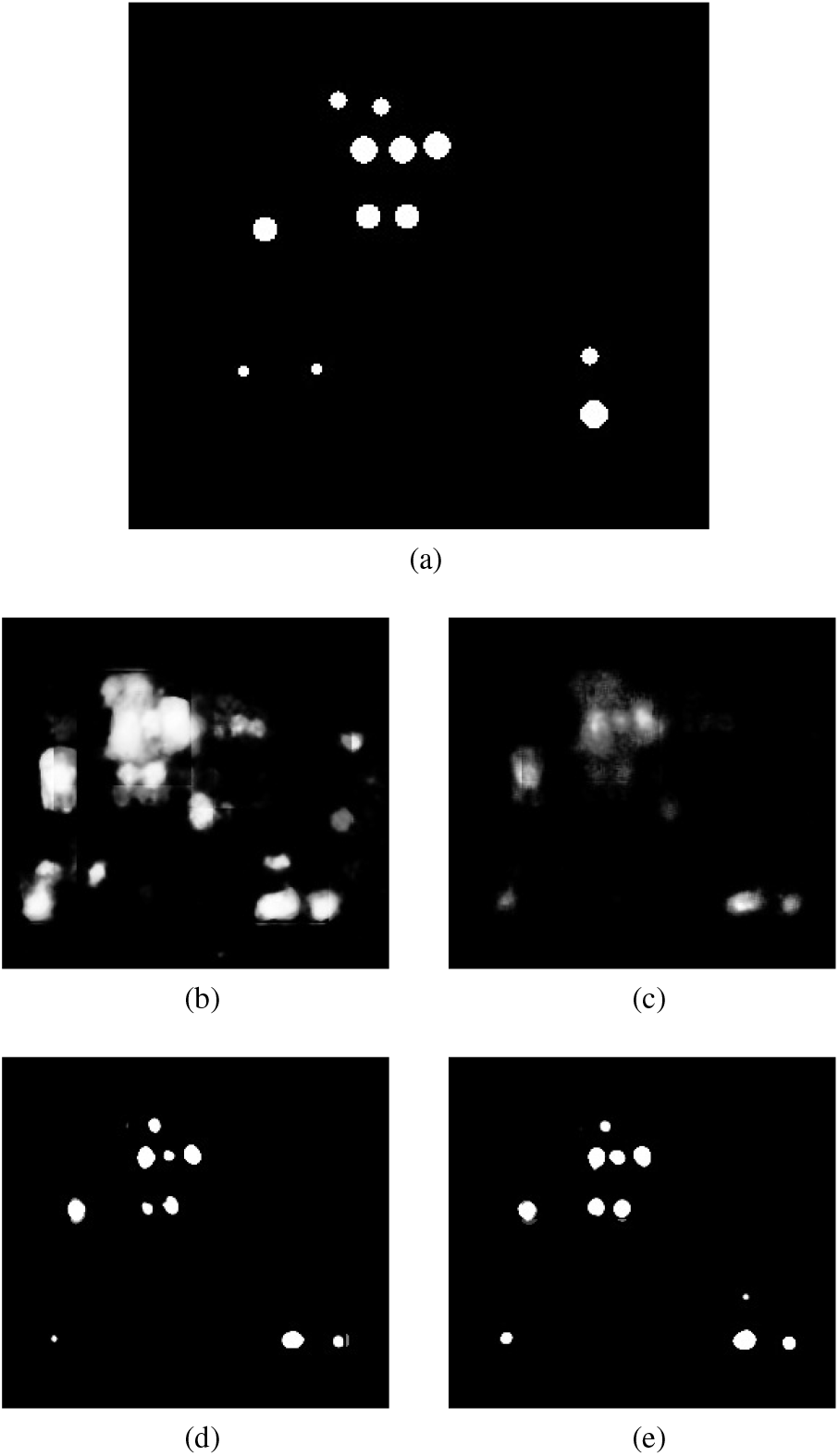
Comparison of segmentation results produced by using different loss functions. (a) shows the ground truth of the input volume labelled regions, network outputs are then shown using the following loss functions: (b) WCE (Weighted Cross Entropy), (c) BCE (Binary Cross Entropy), (d) Dice and (e) GDL (Generalized Dice Loss).

The GDL loss function was proposed by Crum et al. [21] to evaluate segmentation performance using a single score. Sudre et al. [20] then proposed to use this score as a loss function to train deep neural networks to perform segmentation on highly imbalanced data, and is represented as:

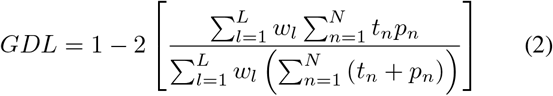

where, *t* is the reference ground truth volume, *p* is the predicted probalistic map and *w* is the label specific weight. The contribution of each label to the GDL weight *w* is determined by the inverse of that label’s volume. This therefore reduces the correlation between the Dice score and the region size. *w* is represented by:

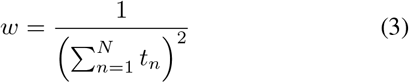

## V. RESULTS

As explained in Section IV-A, the confocal dataset is composed of five time-lapse sequences of five different roots, which contain a total of up to 142 individual time points. Out of these five sequences, four (containing 132 time points total) are used for training and validation while the last (containing six time points) is used for evaluating final performance of the network. At no time during training is any volume from the last test sequence introduced to the network, ensuring adequate separation between training and testing, and allowing a “real world” demonstration of the network as if run on a newly-captured sequence in a subsequent biological experiment. In an effort to visualise the 3D output of our network for this paper, a 2D slice is taken from each volume along with its respective slice in the input volume and ground truth. This representation is shown in Figure 5. Using this the reader should be able to get a qualitative feel for network performance, but do note the true output is a 3D labeling of the data.

**Figure 5:**
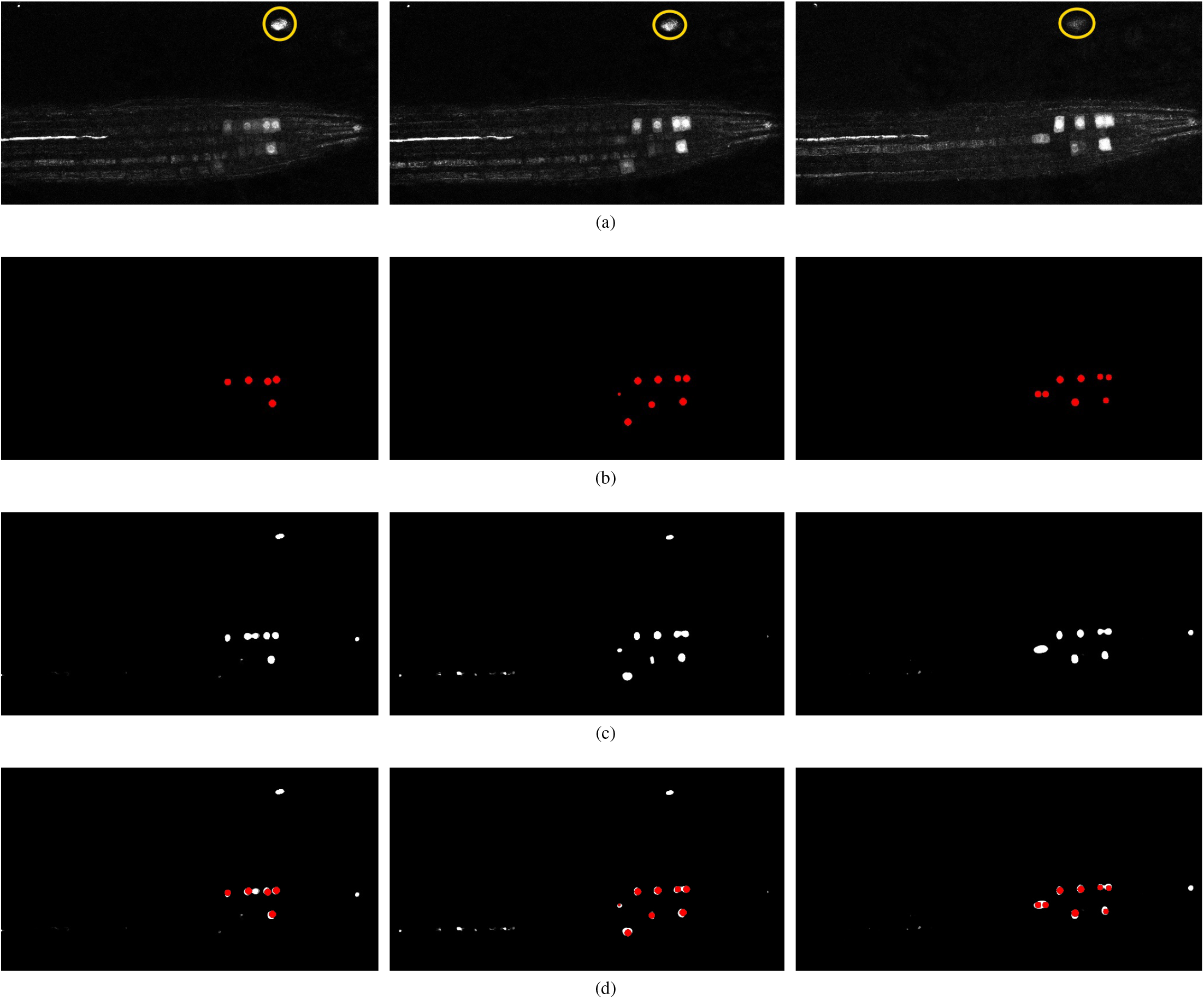
Representation of network output over different time points of the test timeseries (time points 1, 2, and 3), and comparison against the ground truth and the input data. (a) Represents a 2D slice from the input volume (note, the yellow circle indicates an artefact feature referred to in the main text), (b) represents the generated ground truth for this input volume, (c) represents the network output and (d) shows the overlay of the ground truth (red) on top of the network output (white). If comparing to numerical results, note that this is a 2D section of a 3D slice – more nuclei are likely present.

Despite limited training data, the challenge of working with the 3D data, and the artefacts present in the confocal images, the results produced by the network look encouraging. The overlay of the ground truth over the output (See Figure 5(d)) shows that our network has detected almost all of the nuclei markers present in the input volumes along with some false positives; however, comparing these false positives with their respective input volumes shows that the network is arguably justified in detecting these events, as they are located on feasible artefacts in the image (represented as those enclosed within the yellow markers in Figure 5(a)). This clutter presents itself as a feasible marker; however, its location in the image means a biologist would discount it, demonstrating one of the downsides of a patch-based approach – the lack of spatial context. This is considered more in the discussion section. In some places, the markers are in very close proximity. This is likely a common occurrence in this and other biological datasets, as the markers will tend to cluster at sites of similar biological action – in this case, cell division. It can be seen in e.g. Figure 5(c) centre panel that markers in close proximity are sometimes joined, producing a single output volume where there should be two distinguishable volumes in reality. Due to the shape of the regions, however, this error can be easily fixed in a post processing stage using morphological operations, which will be explained in Section V-B.

### A. Quantitative evaluation protocol

Numerical evaluation of our network’s performance is carried out by determining the F-measure value, which is defined as the harmonic mean of precision and recall, and also referred to as the F1 Score. Precision represents the ability of the network not to classify a sample as positive when it is actually negative, and is determined using:

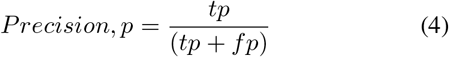

Conversely, the recall value is the ability of the network to detect all of the positive samples present in the volume and is determined using:

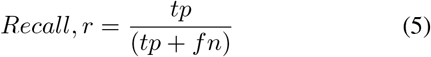

where, *tp* is the number of true positives, *fp* is the number of false positives and *fn* is the number of false negatives. The F-measure is a combination of these values, giving an overall measure of performance and is determined as:

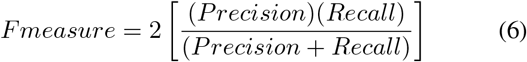

It is important to determine the criteria for what can be considered a correct detection, and to distinguish between a true positive, false positive and false negative. This was accomplished by using a count and distance measure, where a simple Euclidean distance is determined between the centre of a detected nucleus and its closest ground truth nucleus. If this distance is less than a set threshold then it is considered as a true positive, otherwise it is considered as a false positive. Any nuclei that the network fails to detect are counted as false negatives. The value of this threshold was determined empirically and it was observed that a distance threshold of 5 pixels gave the best representation of the network’s performance. In practice this value will be related to the size of the biological feature being marked, the zoom of the microscope, and the calibration scale of the image. Table I reports the F-measure performance of each volumetric test sample given to the network.

**Table I.** Evaluation of network using precision, recall and f-measure, calculated over the 6 timepoints in the test timeseries.

| Volumetric Time Stamp | Nuclei Reported in Ground Truth | Nuclei Predicted in Output | True Positives | False Positives | False Negatives | Precision | Recall | F-Measure |
| --- | --- | --- | --- | --- | --- | --- | --- | --- |
| 1 | 5 | 8 | 5 | 3 | 0 | 0.63 | 1.0 | 0.77 |
| 2 | 11 | 9 | 8 | 1 | 2 | 0.88 | 0.8 | 0.84 |
| 3 | 9 | 9 | 8 | 1 | 0 | 0.88 | 1.0 | 0.94 |
| 4 | 6 | 6 | 5 | 1 | 0 | 0.83 | 1.0 | 0.90 |
| 5 | 3 | 6 | 3 | 3 | 0 | 0.50 | 1.0 | 0.66 |
| 6 | 2 | 5 | 2 | 3 | 0 | 0.40 | 1.0 | 0.57 |

### B. Discussion

It can be seen from the reported results that our network is producing encouraging results even for difficult volumetric samples, particularly given the small number of training instances used. In practice, the network allows cell division markers to be identified, located and, if necessary, counted within a volumetric dataset. The network continues to produce some false positives, however, predicting a marker where there should not be one. Upon further investigation, we found that a false positive reported in almost every sample is actually an artefact (marked in Figure 5(a)) present in the test input volume throughout the time sequence. This artefact resembles a nucleus feature in appearance and is therefore reported as one by the network. Figure 7 shows the presence of this artifact, which can be seen in close-up as the bright spots at the top of the volume. Since this lies outside our pre-determined distance threshold it is thereby reported as a false positive. Such an artefact is easy to throw away by visual inspection because of the context – the location is outside of the root, and so would be dismissed by a biologist. This highlights a downfall of such networks, that context can be lost in process of dividing the image into overlapping patches. It may be possible to build context into the network (indeed this is discussed in the Conclusion section), but here we test a simpler method to prevent it’s detection in the first place.

The underlying problem here is one inherent in machine learning – the training data did not include any such samples as negative ground truth. It is only present in the time-series that was randomly selected for testing the system. Had there been more training samples with such artefacts from which the network could learn, then the network could learn to ignore them. To test this hypothesis, our network was re-trained from scratch on a much smaller and re-shuffled dataset where majority of the artefact stained samples were included into the training pool. But this caused a large imbalance of data as there were very few stained samples compared to the non-stained ones. To rectify this, the training/validation pool was reduced from 132 time points to around 50 time points, out of which 9 volumes had the artefact staining whereas the remaining 41 did not. 2 artefact stained time points were left out for the purpose of testing the network. And as expected the system successfully learned to ignore such artefacts and only reported and segmented the actual nuclei. The output of this re-trained network can be seen in Figure 6, here the artefact is no longer reported even though it is very prominent in the input volume, demonstrating the importance of a suitable training set.

**Figure 6:**
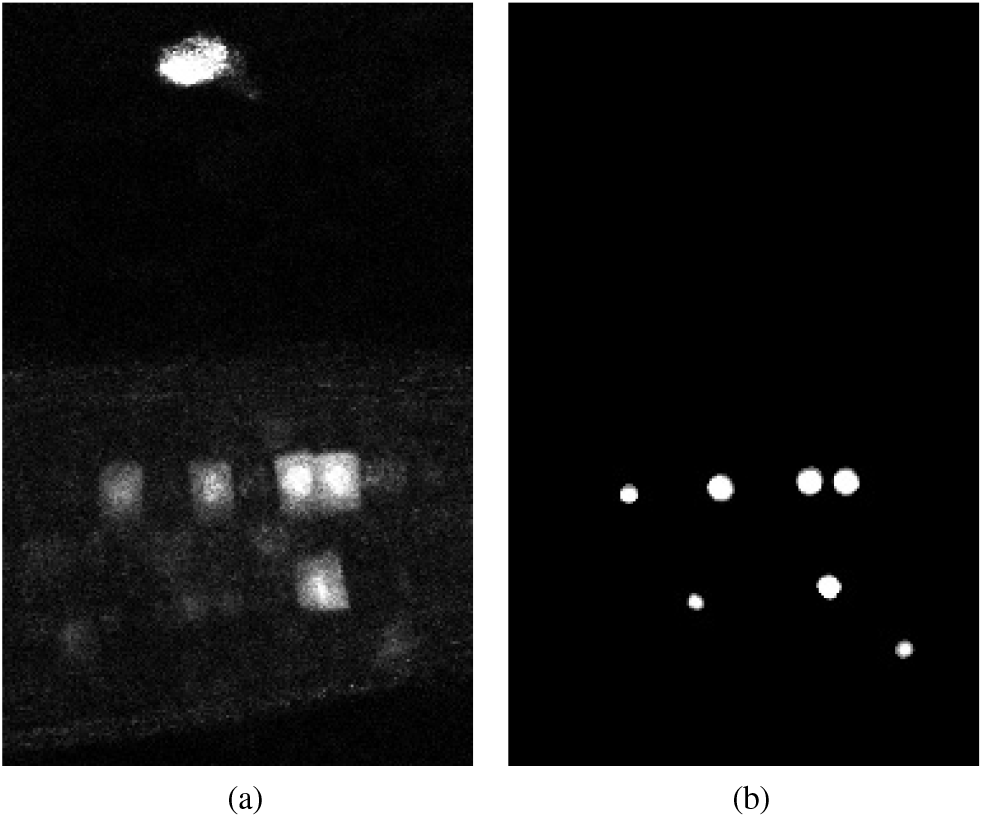
Representation of the output volume (b) for input data (a) produced by the network trained on the re-shuffled dataset, feature a wider variety of training data. Note the artefact outside the root, at the top of the image, is no longer labelled on the output.

**Figure 7:**
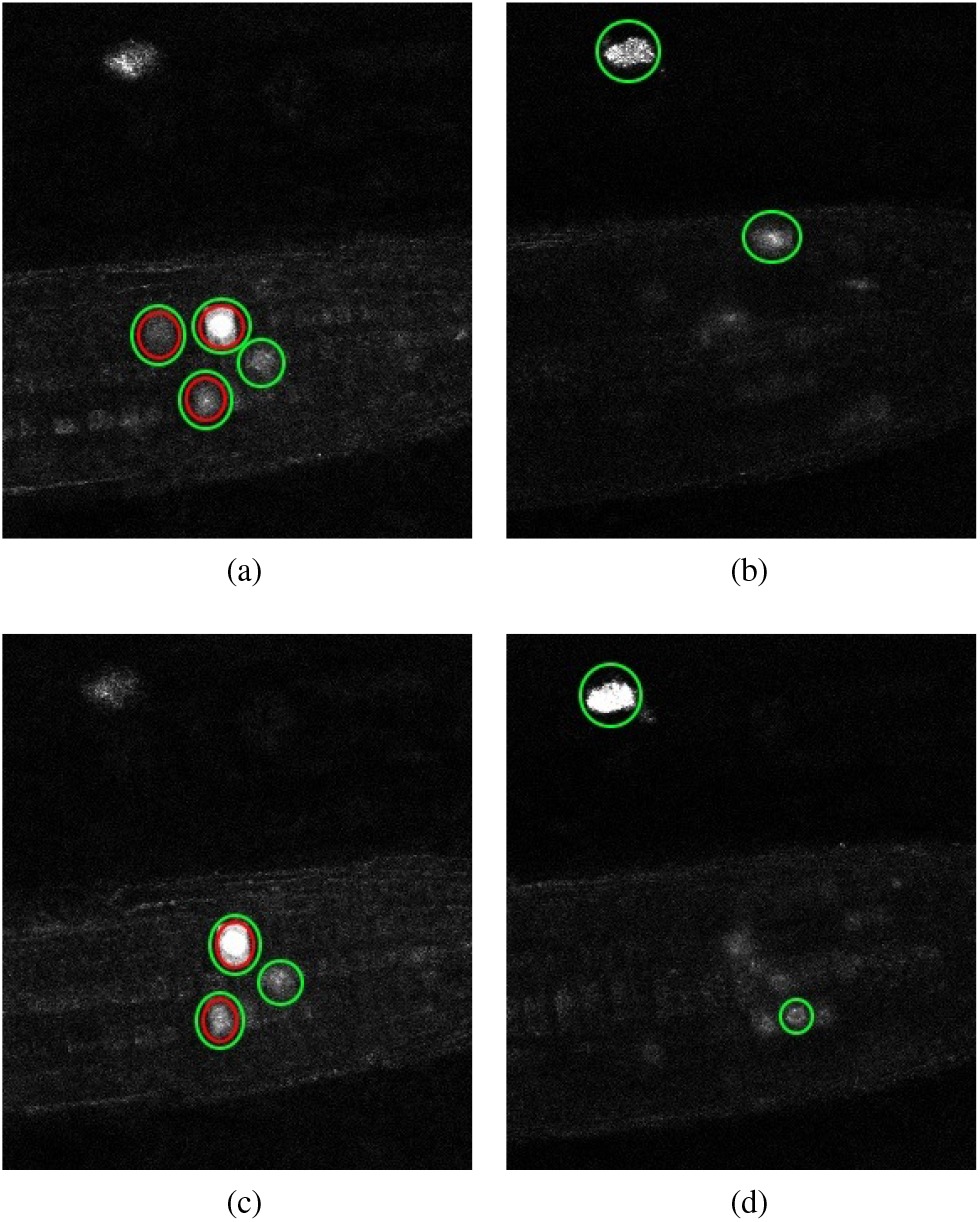
Error cases : artefact detection and ambiguous annotation. Representation of the nuclei detected by our network and those reported in the ground truth. Individual z-slices showing prominent detections taken from test volume 5 are shown in (a) and (b) while that from test volume 6 are shown in (c) and (d). The nuclei enclosed in the green circles are the ones detected by the network and those enclosed in the red circles are the ones reported by the ground truth.

Another error case highlights the sensitivity of such deep learning approaches to annotations themselves. The network labels twice as many markers in volume 5 than are present in the ground truth. Examining the input volume for this time stamp shows why the network may report these locations as viable predictions of the marker. Figure 7 (a) and (b) shows example z-slices from the input in question.

Similarly, the network is reporting 5 nuclei present in volume 6 whereas only 2 are present according to the ground truth. As for test volume 6, Figure 7 (c) and (d) shows different z-slices from this volume where the nuclei have been detected. Green circles represent the network predictions whereas red circles represent the ground truth. As can be seen, the network detections seem justified but since they are not marked as present in the ground truth, these detections are reported as false positives. In the end, it would be left to the biologist to determine whether a detection by the network is a true positive or a false positive; it is likely a judgement call.

As shown in Figure 5, the network struggles to completely distinguish between nuclei that are closely located to each other, but is able to detect two entities that are joined (see Figure 8(a)). Despite the fact that point annotations are used, giving the regions of interest volumes means they can become connected regions as depicted in this figure. However, the shape they acquire when joined is amenable to further processing to distinguish the two regions. This allows us to make use of a combination of morphological operations to clean up the network output and further improve the result. Figure 8(a) shows a 2D slice taken from an output volume that has detected two nuclei as a single connected component; we will now use this volume to explain the post-processing process.

**Figure 8:**
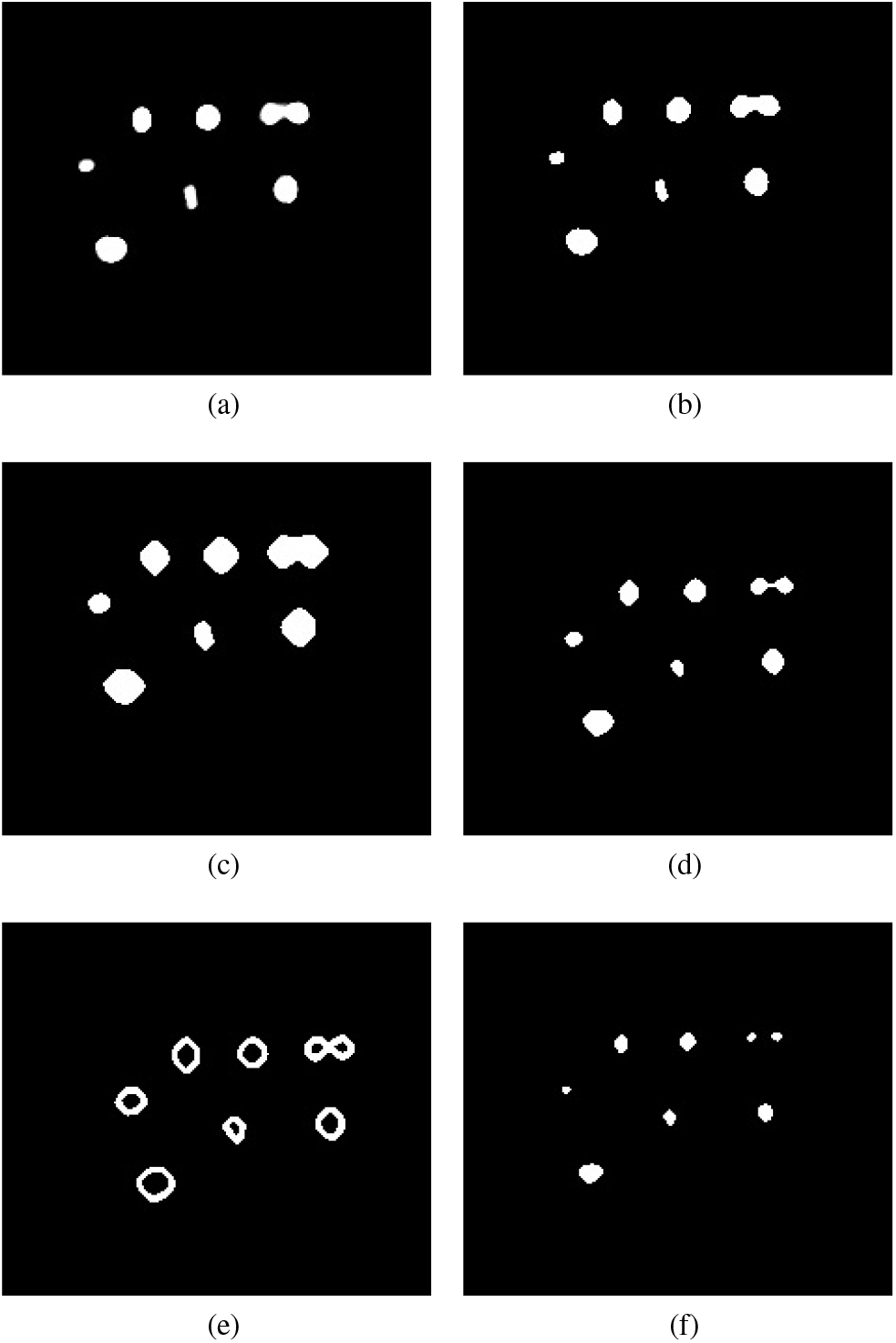
Steps in the post-processing process to distinguish incorrectly joined neighbouring predictions. (a) the predicted output from the network (b) morphological opening applied to remove unwanted noise (c) morphological dilation marks the background pixels (d) distance transform marks the fore-ground pixels (e) subtraction of foreground from background to get the boundary region of connecting nuclei(f) watershed segmentation produces the final result (images cropped to area of interest for clarity).

To begin with, the image quality of the output produced by the network is first improved by removing any unwanted noise and artefacts. This is achieved via morphological opening using a sphere structuring element. The output of this stage is shown in Figure 8(b). Having cleaned up the volume, we may be confident that the region near the centre of the connected components are the foreground and the region further away from the centre are part of the background. The only region that we are unsure of is the boundary region that connects the two nuclei. To mark this boundary region we need to know which part of the connected components are background for certain, and which part of the connected components are foreground. The “sure background” region is obtained by morphological dilation of the volume using the same structuring element, the output of this step marks our “sure background” and is shown in Figure 8(c). Distance transform is then applied on the dilated “sure background” volume, which is then thresholded to get the “sure foregorund” region, as shown in Figure 8(d). Now that we know for certain which areas represent background and which areas represent foreground we can identify the boundary region by subtracting the “sure foreground” volume from the “sure background” volume. The output of this subtraction is shown in Figure 8(e). Having identified all the regions in the volume, the nuclei can be marked as areas of interest while the boundary regions can be marked as area’s to ignore. Finally, watershed segmentation can make use of these marked areas to produce the final output that separates any connected nuclei at the boundaries, as can be seen in Figure 8(f).

## VI. CONCLUSION

We have presented a volumetric deep learning approach which is able to segment labelled nuclei markers in confocal time series datasets with only a limited number of training instances. Training has been carried out in time series datasets isolated from the test data, providing as realistic a use case as possible.

The segmentation was carried out on discrete time points. A future development of this work will actively try to segment time-based events such as cell divisions, linking features over time perhaps using a tracking approach. This process will be performed on light-sheet data (versus confocal), as light-sheet microscopy is able to capture many more volumes both in frequency and period, but the work here will provide a foundation for this development.

The results demonstrated that unless particular care was taken with the training data coverage, it was easy for the network to mistake clutter for desired features, even if the clutter was located outside of the biological root system. The developed network makes use of volumetric patches in order to overcome memory demands on the GPUs. However, results here suggest contextual information could be useful to provide to the network, such that objects outside of biologically relevant areas could be discounted. Previous work has used lower resolution data with more spatial coverage to provide such context, and so such an approach could be utilised here in the future.

Of particular note is that the results were achieved with only a limited amount of simple augmentation. Given the relatively constrained problem we are handling – that of particular markers in a root oriented in the same direction in each image, captured using very similar microscope settings – these augmentations seem to be sufficient. However, there is potential work in developing confocal-microscope specific augmentation. This could include, for example, synthesising noise expected to affect confocal images, as well as simulating the effects of laser attenuation throughout the sample. Such augmentation could be expected to lessen the sensitivity of the training to particular features of the training set, which is even more important where training set size is limited – as is often the case with 3D microscopy due to the overhead in annotating the images, as well as the level of expertise required to do this.

In this paper we have been concerned with finding fluorescent markers which indicate a growth event (in this case cell division) happening as part of the growth of a root. Therefore, when evaluating the approach, we are primarily interested in if we produce a prediction sufficiently close to a ground truth *x, y, z* label (a true positive) rather than a prediction away from one of these labels (a false positive). We also want to make sure ground truth labels are not missed (false negatives). In future work, we can refine the segmentation of the marker region itself, i.e. evaluate the pixel-wise segmentation of the region. Whilst this resulting volume or shape information is not relevant for the marker in use here, it could be important if trying to analyse the shape of a cell as it expands or divides, for example. As a pixel-wise segmentation is already produced by the network, but refined in post processing to a single location in space, the network is already partially capable of generating meaningful 3D shape labels.

## VII. ACKNOWLEDGEMENTS

This work was supported by the Biotechnology and Biological Sciences Research Council [BB/N018575/1].

